# Aging of human endocrine pancreatic cell types is heterogeneous and sex-specific

**DOI:** 10.1101/729541

**Authors:** Rafael Arrojo e Drigo, Galina Erikson, Swati Tyagi, Juliana Capitanio, James Lyon, Aliya F Spigelman, Austin Bautista, Jocelyn E Manning Fox, Max Shokhirev, Patrick E. MacDonald, Martin W. Hetzer

## Abstract

The human endocrine pancreas must regulate glucose homeostasis throughout the human lifespan, which is generally decades. We performed meta-analysis of single-cell, RNA-sequencing datasets derived from 36 individuals, as well as functional analyses, to characterize age-associated changes to the major endocrine pancreatic cell types. Increasing age was associated with shifts in pancreatic alpha and beta cell identity and loss of nuclear integrity in non-diabetic humans. In non-diabetic individuals ≥ 50 years old, 80% of their beta cells exhibited a transcriptional signature similar to cells from type-2 diabetic (T2D) donors. Surprisingly, ∼5% of beta cells from T2D donors retained a youthful, N.D. transcriptional profile. Furthermore, beta cell function was reduced by 50% during aging in men but not women, which may explain sex-associated differences in diabetes etiology. These analyses reveal that aging of the human endocrine pancreas is sex- and cell-type specific.

## Introduction

Aging is characterized by the functional deterioration of proteins, organelles, cells and tissues (Ori et al., 2015; Petersen et al., 2003; Taylor and Dillin, 2011). These progressive declines are associated with a higher incidence of neurodegenerative, cardiovascular, and metabolic diseases in aged humans. Aging increases the risk of developing type 2 diabetes (T2D), a disease that further increases the risk of cardiovascular complications and death (Kalyani et al., 2017). T2D is primarily caused by the functional decline and/or loss of pancreatic beta cells and loss of peripheral insulin signaling, which results in impaired glucose homeostasis (Doria et al., 2008; Gutiérrez et al., 2017). Beta cells are located in the islets of Langerhans and secrete insulin in response to rising glucose levels (Cabrera et al., 2006). Beta cell function is regulated by circulating and paracrine factors secreted by neighboring alpha cells and delta cells, as well as the autonomic nervous system (Arrojo e Drigo et al., 2015).

The major endocrine cells of the human islet, namely the alpha, beta, and delta cells, are thought to be largely long-lived cells that can be as old as post-mitotic cortical neurons (Arrojo e Drigo et al., 2019; Cnop et al., 2010). Like neurons, human islet cells must maintain functional integrity for decades if they are to sustain glucose homeostasis throughout the human lifespan. In the absence of cell division, long-lived cells suffer long-term exposure to toxic metabolites and the accumulation of amyloid protein aggregates, which result from the loss of cellular repair processes and protein quality control mechanisms (Taylor and Dillin, 2011). Therefore, understanding how functionality of these cells is maintained and lost during aging is important for understanding the basic biology of the pancreas and T2D pathophysiology. Notably, previous studies have shown that islet beta cells are a functionally heterogeneous population of cells, displaying different activity patterns in response to glucose stimulation (Van Schravendijk et al., 1992). Recent advances in light microscopy and single-cell transcriptomics have confirmed this result, and further shown that such functional heterogeneity may be driven by distinct transcriptional states that give rise to beta cell “sub-types” (Camunas-Soler et al., 2019; Dorrell et al., 2016; Johnston et al., 2016; Segerstolpe et al., 2016; Xin et al., 2016, 2018).

While these studies have revealed important aspects of mostly beta cell biology, the impact of aging on beta cells and other human islet cell types remains largely unknown. Data derived from humans islets indicate that aging may be associated with: 1) changes in the overall chromatin state of beta cells (Avrahami et al., 2015), 2) a decline in beta cell proliferation (Perl et al., 2010), and 3) increased transcriptional noise in both alpha and beta cells (Enge et al., 2017). While adult beta cells can secrete more insulin than those of children and adolescents, likely due to their advanced maturation state (Avrahami et al., 2015), it is still unclear whether beta cell function continues to evolve throughout adulthood. For example, using isolated islets, one study found no differences in insulin secretion throughout adulthood (Almaça et al., 2014), whereas others observed a decrease in beta cell calcium dynamics and insulin secretion in humans greater than 40 years of age (Westacott et al., 2017). Furthermore, sex-specific differences in glucose homeostasis and DNA methylation patterns suggest that islet cells from men and women could have different transcriptional and functional profiles (Basu et al., 2006; Hall et al., 2014) and this could underlie sex differences in T2D etiology (Logue et al., 2011).

Here, we describe the effects of aging and sex differences on the transcriptional phenotype of major islet cells in 26 non-diabetic (N.D.) donors spanning several decades of life, and 10 T2D donors. We integrated three recently published RNA-sequencing (RNAseq) datasets derived from the human pancreas (Enge et al., 2017; Segerstolpe et al., 2016; Xin et al., 2016) and performed meta-analysis followed by analysis using the Pagoda2 and Monocle v2.4 platforms. We identified robust transcriptional profiles for sub-populations of alpha, beta, and delta cells, and showed that 75% of aged beta cells have a transcriptional signature similar to that of beta cells from T2D donors. T2D beta cells were heterogeneous, with different beta cell subpopulations exhibiting progressive losses in cell identity and function. Strikingly, ∼5% of T2D beta cells maintained a phenotype associated with optimal beta cell function, which was typically seen in young N.D. adults. Finally, aging was associated with impaired beta cell function in men but not women. This phenotype was reflected in transcriptional profiles that indicated a loss of beta cell identity and function only in male beta cells. Together, our data highlight how aging and sex contribute to the physiology of different long-lived populations of cells within the human pancreas in both health and disease.

## Results

### Meta-analysis of three single-cell RNAseq datasets derived from the human pancreas

Aging of post mitotic tissues is generally associated with declines in the function, cell identity, and/or protein homeostasis of long-lived cells (D’Angelo et al., 2009; Taylor and Dillin, 2011). Alpha and beta cells are both post-mitotic cells that exhibit limited amounts of turnover during an organism’s lifetime (Arrojo e Drigo et al., 2019; Cnop et al., 2010), which in humans can last several decades. As such, we hypothesized that the cellular identity and/or function of alpha and beta cells would decline with age in N.D. adult humans. To test this hypothesis, we combined the datasets from three recent human islet single-cell RNAseq studies and performed meta-analysis. The combined dataset contained data from 36 donors (26 N.D. subjects, ages 1 month to 68 years; and 10 patients with T2D) (Figure 1A, Table S1 and methods) (Enge et al., 2017; Segerstolpe et al., 2016; Xin et al., 2016). We chose this approach because each study alone does not contain a sufficient number of donors to cover the human lifespan, thus making it difficult to study islet cell aging in adults.

**Figure 1.**
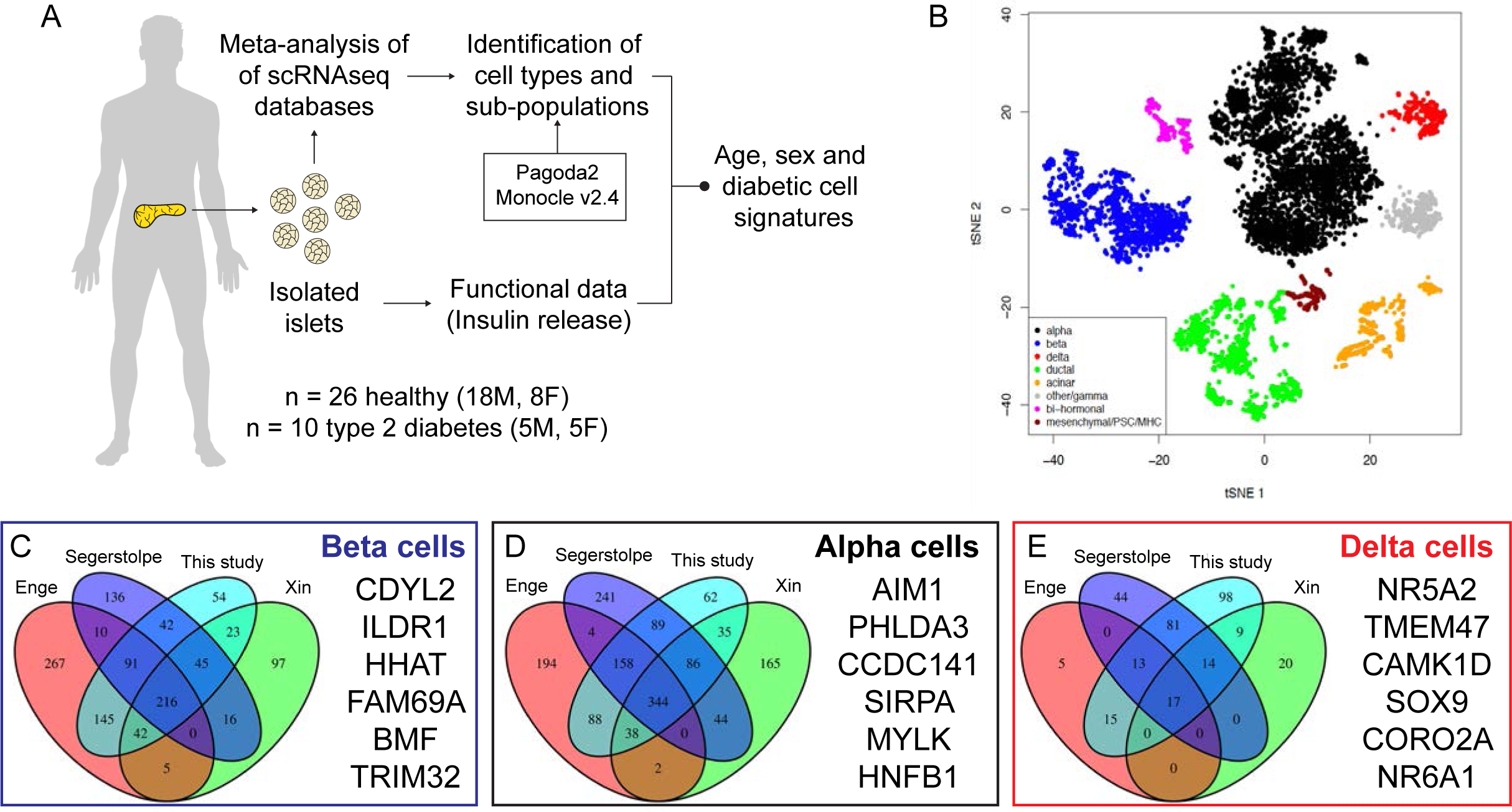
Meta-analysis of single islet cell RNA seq datasets. **(A)** Illustration describing this study’s schematics. **(B)** tSNE clustering of the Combat-corrected single cell RNAseq data labelled with the identified cell types. In **(C)** to **(E)**, Venn diagrams showing the overlap in the number of differently expressed genes identified in beta cells, alpha cells and delta cells (respectively) by in each individual study (Enge et al, Segerstolpe et al and Xin et al) versus our meta-analysis approach (this study).

We first used ComBAT to re-align, filter, and batch correct the different single-cell datasets, resulting in a single dataset that contained ∼6500 pancreatic cells from the 36 donors (Figure S1A-B). Next, we identified all major pancreatic endocrine and exocrine cell types based on relative expression levels of cell type-specific marker genes (Figure S1C-I, Table S2, methods), which were highly variable in alpha, beta, and delta cells (Figure S1J). Cell compositions were variable across the different donors (Figure S1K), and analysis of each individual dataset revealed limited overlap between genes expressed by alpha, beta, and delta cells (Figure S1L). Our meta-analysis identified new genes that were specific for beta (n = 54) and alpha (n = 62) cells, but the largest number of new marker genes were identified for delta cells (n = 98), which included expression of the stem cell markers *NR5A2, NR6A1* and *SOX9* in this rare islet cell type (Figure 1C-E). These results show that our meta-analysis approach effectively integrated different single-cell RNAseq datasets, removed study-specific data biases, and identified new cell type-specific genes.

### Age-dependent loss of alpha and beta cell identity in N.D. humans

To determine the effect of aging on adult alpha and beta cells, we divided the sample cohort into three age ranges: 1) young, which were from donors less than 6 years old (y.o.), 2) adults, which were from donors 20–48 y.o., and 3) aged adults, which were from donors 54–68 y.o. These age ranges were selected for two main reasons: *i)* to separate very young samples, which are still undergoing postnatal organ development, from more differentiated and mature cells within the adult pancreas, and *ii)* because the incidence of diabetes nearly doubles in humans 45 y.o. or older (Table 1, National Diabetes Statistics Report, 2017, Center for Disease Control).

We performed differential gene expression and pathway enrichment analysis for the bulk alpha and beta cell populations in adults versus aged adults, and identified 149 genes (n = 108 up, n = 41 down, FDR ≤ 1%) and1325 genes (n = 998 up, n = 327 down, FDR ≤ 1%) that were differentially expressed in aged beta and alpha cells, respectively (Table S2-3). Age-related changes in these cells were defined by a decline in the expression of genes involved in cell identity and exocytosis, and an up-regulation of genes characteristic of other endocrine cell lineages (Figure S1M, Table S3-4). Both cell types in aged adults up-regulated genes involved in ribosome biogenesis and translation pathways, which are markers of aging in human cells (Buchwalter and Hetzer, 2017). However, we did not observe changes in the islet aging markers, IGF1R or p16/CDKN2A (Table S3-4, (Aguayo-Mazzucato et al., 2017; Helman et al., 2016)). Together, these data provide evidence that the cell identity of most alpha and beta cells in aged N.D. humans is compromised (Figure S1M, Tables S3-4).

### Identification of alpha and beta cell subpopulations

Alpha and beta cells are transcriptionally and functionally heterogeneous (Camunas-Soler et al., 2019; Dorrell et al., 2016; Gutierrez et al., 2018; Johnston et al., 2016; Van Schravendijk et al., 1992; Segerstolpe et al., 2016; Xin et al., 2018) and a recent study indicated that sub-populations of these cells may change with age (Wang et al., 2016). Therefore, we used the Pagoda2 pipeline (Fan et al., 2016), followed by pathway enrichment analysis, to identify sub-populations of alpha and beta cells, with the goal of assessing the impact of aging on cell heterogeneity.

This approach identified three main beta cell states (β-state 1, 2, and 3), each with a distinct gene expression profile (Figure 2A, S2A and TableS5). Of note, a fourth state was identified (β-state 4), however it was comprised of cells derived primarily from a single donor (a 21 y.o. male from (Enge et al., 2017)) and therefore was not considered further for analysis (Figure S2A, Table S5). Differential gene expression analysis revealed that cells in β-state 1 expressed the highest levels of genes involved in beta cell identity (*INS, PDX1, NKX6-1, MAFA*, and *RBP4*), insulin secretion, glucose metabolism, and mitochondrial function (Figure 2A, Table S5). These cells also expressed high levels of *ID1* and *ID3*, and thus were similar to two beta cell sub-populations described in one of the datasets used in our-meta-analysis (Segerstolpe et al., 2016). Cells in β-state 2 expressed genes involved in p53-dependent regulation of transcription and chromatin modification, Wnt signaling (*TCF7L2*), and the insulin-IGF1 receptor signaling pathway (*INSR, IGF1R*), as well as lower levels of the beta cell heterogeneity marker CD9 (Dorrell et al., 2016)(Figure 2A, Table S5). In contrast, cells in β-state 3 were enriched in genes involved in ribosomal biogenesis, mRNA translation, and assembly of the primary cilium (Figure 2A, Table S5) Like β-state 2, cells in β-state 3 expressed genes involved in beta cell identity and function at lower levels (Figure 2A, Table S5). In addition, these cells had higher levels of *TMEM27, SYT16*, and *ASB9* expression, which is consistent with previous data gathered from sequencing of bulk beta cells from juvenile donors (Arda et al., 2016). Neither of these groups expressed “beta cell disallowed genes”, which in mice are associated with loss of beta cell identity and function (Figure S2C)(Pullen et al., 2010)).

**Figure 2.**
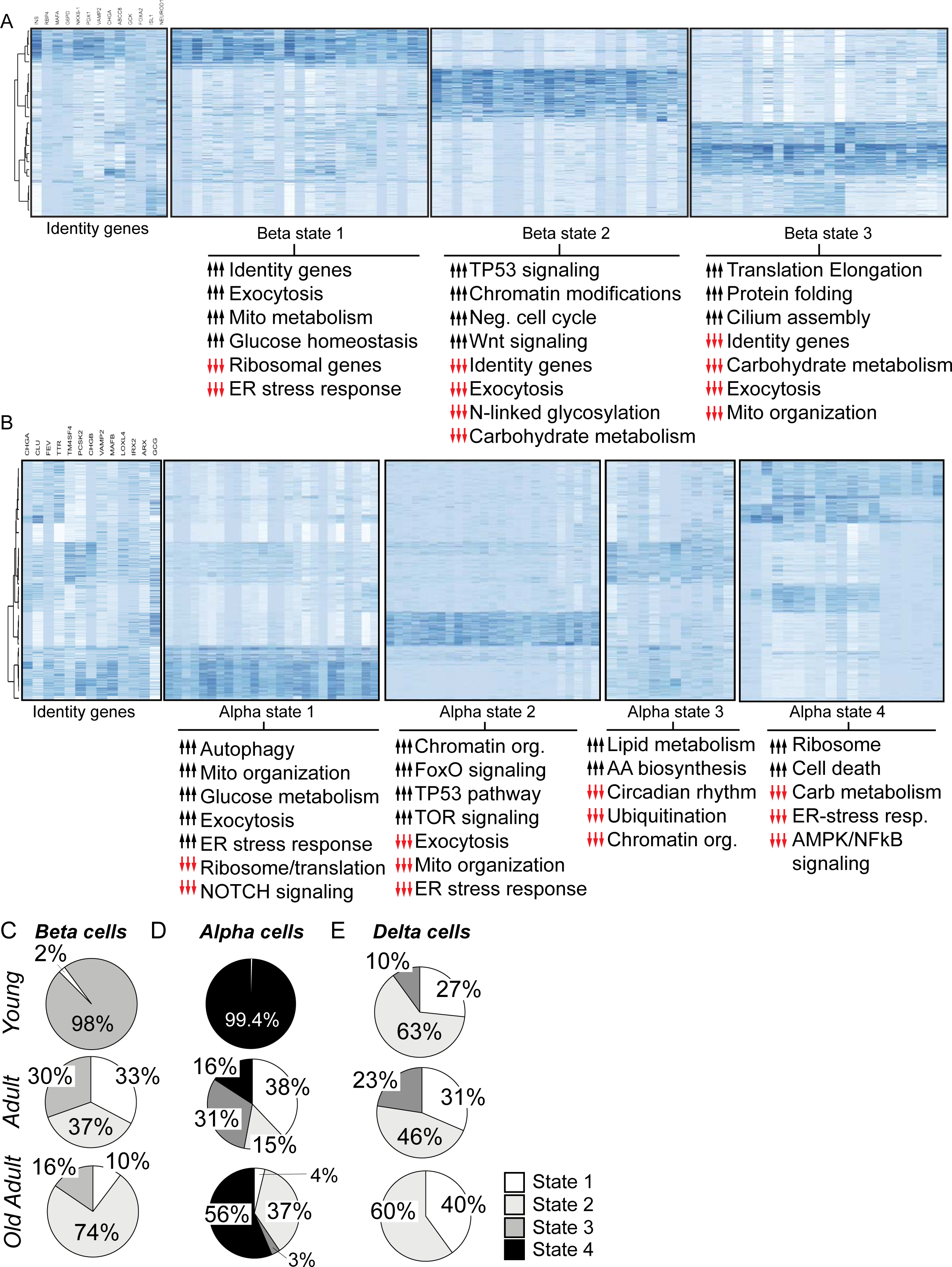
Aging leads to changes in the composition of alpha and beta cell populations. Identification of different types of alpha cell and beta cells based on their transcriptional states. In **(A)** and **(B)**, hierarchical clustering analysis of beta cells (A) and alpha cells (B) from N.D. donors. Heatmaps show the normalized expression of cell identity marker genes (left panels, gene names listed on top) and representative genes differentially expressed in each alpha and beta cell state identified. Below each heatmap, pathway enrichment analysis (Metascape) for each cell state. Representative genes were chosen based on their Z-scores calculated within Pagoda2. Black arrows and red arrows indicate pathways that are positively and negatively regulated, respectively. For a full list of differentially expressed genes in each cell state, please refer to Tables S4-5. **(C-E)** Relative distribution of alpha cells (C), beta cells (D) and delta cells into each state identified with Pagoda2 and as a function of age. Samples were divided into young (<6 y.o.), adults (20-to-48y.o.) and old adults (54-to-68y.o.). **(F)** Glucose-stimulated insulin secretion (GSIS) assay performed on islets isolated from adults ages 18-to-48 y.o. (n=53 donors) and old adults aged 50.y.o. or older (n=80 donors). Islets were exposed to 1mM and 16.7mM and insulin secretion is displayed as a relative % of the total islet insulin content. Error bars represent the 95% confidence interval of the data, where **p=0.0026.

In alpha cells, we identified six cellular states (α-states 1–6), however states 5 and 6 were comprised of cells derived primarily from single donors (a 21 y.o. male and a 38 y.o. female, respectively and both were from the same study (Enge et al., 2017)) and therefore were not considered further (Figure S2D). In contrast to beta cells, the expression of alpha cell marker genes was heterogeneously distributed across these six cell states (Figure 2B, S2D). Cells in α-state 1 had lower levels of GCG expression, and high levels of expression for genes involved in alpha cell identity (*MAFB, ARX*, and *FEV*), glucose metabolism, exocytosis, ER-stress response, and mitochondrial organization (Figure 2B, Table S6). A similar transcriptional signature has been associated with higher Na+ currents and increased cell size (Camunas-Soler et al., 2019), which may indicate cells with increased excitability. Cells in α-state 2 expressed genes involved in the FoxO signaling pathway, chromatin re-organization, mTOR signaling; α-state 3 was characterized by the expression of *TM4SF4*, as well as lipid and amino-acid metabolism genes; α-state 4 was defined by the expression of ribosome and apoptosis genes (Figure 1B, Table S5). These data indicate that our meta-analysis preserved the results derived from its underlying dataset and, most importantly, identified new cellular states and defined their molecular signatures. This included the finding that alpha cell sub-populations may have different metabolic needs (carbohydrates (α-state 1) versus amino-acids and/or lipids (α-state 3)) (Figure 2B).

### Age-related changes in alpha and beta cell sub-populations impair cell function

Aging is associated with a higher incidence of T2D in humans (Basu et al., 2003), which is explained by a decline in mechanisms for maintaining glucose homeostasis (Basu et al., 2003; Petersen et al., 2003). We hypothesized that this phenotype was associated with age-dependent changes to the composition of alpha and beta cell states (Figure 2).

To test this hypothesis, we determined which donor groups were represented in the identified alpha and beta cell states. We found that 98% of beta cells from young donors were in β-state 3, and the rest were in β-state 1 (Figure 2C). In contrast, adult beta cells were equally distributed in states 1–3, whereas 74% of beta cells from aged donors were in β-state 2 (Figure 2C). A similar pattern was observed for young, adult and aged alpha cells (Figure 2D). We compared this aging phenotype with that of delta cells, which are remarkably long-lived and regulate the secretory function of alpha and beta cells (Arrojo e Drigo et al., 2019; Hauge-Evans et al., 2009). We identified three different delta cell states (δ-state 1–3), characterized primarily by the differential expression of a small set of genes (Figure S2E, Table S7). Delta cell aging was associated with only small changes in the composition of the delta cell population (Figure 2C). This result was maintained when we re-classified cells into donors aged 20–40 y.o. or ≥ 40 y.o. (Figure S2F). These results indicate that the composition of the delta cell population changes very little from early childhood, and is instead stable for most of the human lifespan. In contrast, the composition of aging alpha and beta cell populations shifts toward cells that express low levels of genes involved in cell identity and glucose metabolism (Figure 2).

These observations led to the hypothesis that aging result in impaired beta cell function, which would contribute to the impaired glucose homeostasis observed in aging humans (Basu et al., 2003). Previous studies have addressed this question and found conflicting results (Almaça et al., 2014; Avrahami et al., 2015; Helman et al., 2016; Westacott et al., 2017). Here, we analyzed data from a glucose-stimulated insulin secretion (GSIS) assay performed on human islets isolated (n=153 isolations) from N.D. donors 18–84 y.o. with normal HbA1c levels and a mean BMI of 27 ± 5.3 (Table S9). Aging was not associated with lower islet insulin content (adults: 45.758 ± 4.347 ug/islet vs old adults: 46.931 ± 3.17 ug/islet, p=0.8240), however aged beta cells secreted ∼30% less insulin when stimulated with high glucose (Figure 2F). These results indicate that the reconfiguration of alpha and beta cell sub-populations, due to changes in their transcriptome, is associated with impaired beta cell insulin secretion, thus likely contributing to age-related declines in glucose homeostasis (Basu et al., 2003).

### Beta cells from aging N.D. donors exhibit features of T2D

Impairments in cell identity and function observed in pancreatic cells derived from aging humans resembled what has been observed in beta cells from *de facto* T2D donors (Talchai et al., 2012). To determine whether dysregulation of beta cell gene expression in aged N.D. donors is similar to that of T2D donors, we used the Pagoda2 pipeline to analyze single beta cell RNAseq data from age-matched N.D. and T2D donors (mean 51.38 ± 9.9 vs 47.86 ± 7.7 y.o., respectively (p = 0.4639)). We identified two cellular states that contained cells from N.D. and T2D donors (Figure S2B-D). One state contained cells with higher expression of genes involved in beta cell function and identity (Table S9), whereas the second state was enriched for cells that expressed genes involved in mRNA catabolism, chromatin organization, nuclear export, and protein degradation (Figure S3B). Strikingly, ∼25% of beta cells from aged N.D. and T2D donors maintained a gene expression signature characteristic of young, N.D. adult beta cells (Figure 2, Tables S4, S9). Thus, a large fraction of beta cells from aged N.D. and T2D donors exhibited impaired transcriptional signatures, but some beta cells seemed to remain functionally intact.

We validated these results using pseudotime ordering (Monocle v2.4, methods)(Qiu et al., 2017). We identified 4 distinct pseudotime branches containing beta cells from N.D. and T2D donors (Figure 3A, S3D-E). Notably, the distribution of cells in the pseudotime tree was independent of donor age and study (Figure S3F, not shown). We determined the transcriptional signature of each branch by performing differential gene expression analysis (Figure S3H-K, Table S10) and observed one pseudotime branch that contained cells from both N.D. and T2D donors (Figure 3A, S3B, black bounding box). Cells in this specific state expressed high levels of genes involved in beta cell identity and function, similarly to younger adult beta cells (Figure 3B, S3D-E, Table S10). These data recapitulate our previous observations with the Pagoda2 algorithm (Figure S3B-C), and support the hypothesis that T2D islets contain beta cells with a transcriptional program similar to that of adult N.D. cells (Figure 2B, 3A-B).

**Figure 3.**
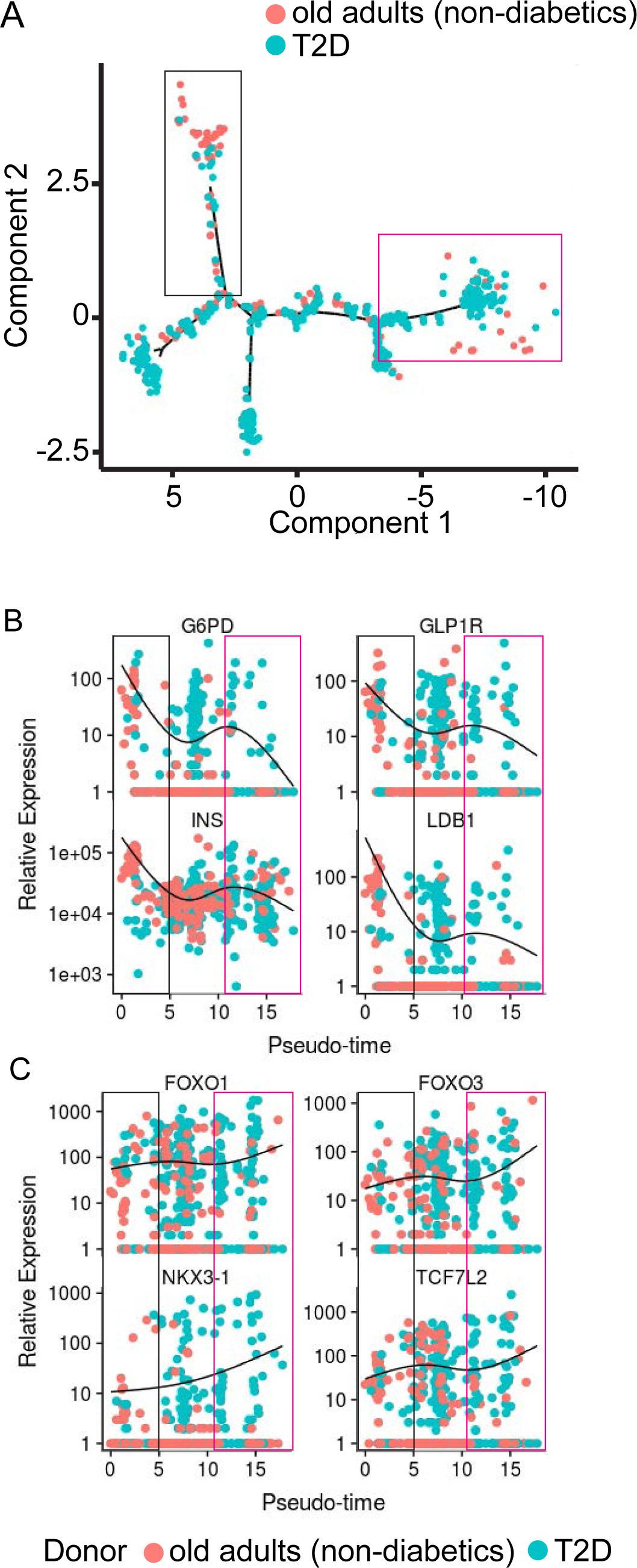
Functional decline of old beta cells is associated with a diabetic-like beta cell transcriptional phenotype. Aging beta cells have a T2D-like phenotype. **(A)** Pseudotime ordering of single beta cell RNAseq data using Monocle (v2.4) of age-matched old N.D. adult donors (red circles) and T2D donors (light green circles). **(B) and (C)**, 2-D pseudotime jitter plots displaying the expression pattern of beta cell identity genes (B) and of genes associated mainly with a T2D beta cell signature **(D)**. The black line in bold indicates the mean expression pattern across the pseudotime tree. In (B-C), black and pink rectangles represent states with high and low expression of beta cell identity genes, respectively.

The remaining cells in the pseudotime tree had transcriptional profiles characterized by step-wise declines in beta cell genes. This pattern had an intermediary branch with ∼25% of cells from aged N.D.s and ∼16% of cells from T2D donors, exhibiting a moderate down-regulation of beta cell identity genes (Figure 3B, S3E, Table S10). The terminal beta cell branch was characterized by high expression of FoxO1/3 genes and the diabetes susceptibility gene *TC7FL2*, in addition to lower levels of expression for glucose metabolism genes and calcium channel genes (Figure 3C, S3G). Strikingly, beta cells in this specific branch displayed a significant decline in the expression of the glucagon-like peptide-1 receptor (*GLP1R*), which mediates the action of GLP-1 on beta cells and enhances glucose-stimulated insulin secretion (Figure 3B, Table S10). These results indicate that T2D islets contain normal beta cells as well as cells characterized by a decline in the expression of cell signature genes (Figure 3A).

### Reconfiguration of the human delta cell population in T2D islets

Recent studies have implicated delta cells in diabetes pathophysiology (Unger and Orci, 2010; Weir et al., 1981; Xin et al., 2016), however the underlying transcriptional signature of diabetic delta cells is largely unknown. To address this question, we used Monocle to analyze data from age-matched N.D. and T2D donors, and identified distinct delta cell sub-populations (Figure 4A). Delta cells from N.D. donors were distributed primarily between two branches, whereas most T2D cells were clustered in a single distant branch (Figure 4A-B). The two main delta cell sub-populations in N.D. islets were distinguished by the expression of MALAT1, as expected from our previous analysis (Figure S2E, Table S11). Differential gene expression analysis of delta cells that expressed high or low levels of MALAT1 (delta^MALAT1high^ and delta^MALAT1low^, respectively) revealed that delta^MALAT1high^ cells expressed lower levels of 697 genes, which included those involved in nuclear and chromatin organization, and glucose metabolism (e.g., LMNA, HDACs 3,5 and G6PD, respectively) (Figure 4C, Table S11). Delta^MALAT1low^ cells expressed high levels of genes associated with mitochondrial metabolism, exocytosis, and protein translation (Figure 4A-C, Table S11). In contrast, T2D delta cells expressed higher levels of lipid metabolism and histone modifications genes, many of which have not been previously associated with diabetes (Figure 4D, Table S11). Notably, diabetic delta cells expressed low levels of genes associated with the delta^MALAT1low^ population (Figure 4C). Together, these data indicate that T2D diabetes leads to declines in the expression of delta cell identity genes and changes the delta cell heterogeneity landscape normally observed in N.D. islets.

**Figure 4.**
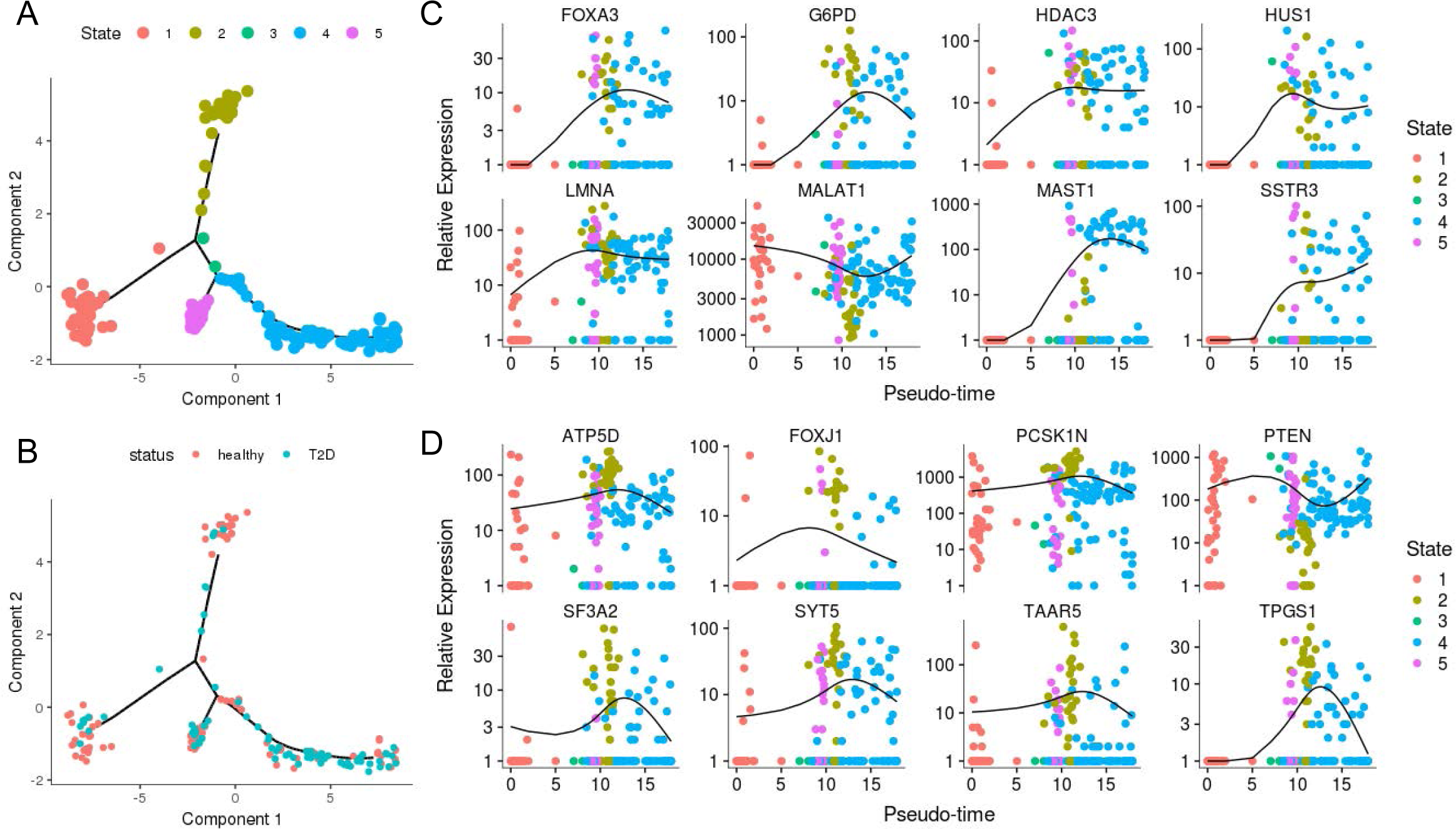
Diabetes leads to a shift in delta cell’s transcriptional phenotype. **(A)** and **(B)**, pseudotime ordering of single delta cell RNAseq data using Monocle (v2.4) of age-matched old N.D. adult and T2D donors. In (A), different states of delta cells are shown and in (B), red and green circles represent cells from N.D. donors and T2D donors, respectively. **(C-D)** 2-D pseudotime jitter plots displaying the expression pattern of differentially expressed genes in delta cell states shown in (A). The black line in bold indicates the mean expression pattern across the pseudotime tree.

### Sex specific differences in alpha and beta cell identity and function

Glucose homeostasis is maintained differently between men and women (Anderwald et al., 2011; Basu et al., 2006; Færch et al., 2010; Gannon et al., 2018). Although both sexes have similar peripheral insulin sensitivity, women have lower levels of endogenous glucose production, lower fasting plasma glucose concentrations, and higher glucose levels ∼2 hours after delivery of an oral bolus of glucose (Anderwald et al., 2011; Basu et al., 2006; Færch et al., 2010; Gannon et al., 2018). While these differences can be partially explained by differences in body mass composition and/or circulating sex hormone levels (Logue et al., 2011; Mauvais-Jarvis et al., 2017; Sicree et al., 2008), little is known about the sex-dependent transcriptomic and functional signatures of human islet cells, and the effect of aging on islet secretory function.

First, we set out to determine whether there are differences between males and females with regard to alpha and beta cell function. We revisited our GSIS experiments (Figure 2F) and separated the data only by sex. Beta cells from adult women secreted 34% less insulin than those from men (Figure 5A, Table S9). No differences in insulin content were observed (Figure S5A, Table S9). Similar insulin release levels have been described in perifusion experiments involving N.D. islets (available from the Human Pancreas Analysis Program (HPAP) (https://hpap.pmacs.upenn.edu/)). Our data also revealed that alpha cells from women secreted less glucagon, despite no differences in islet oxygen consumption or islet cell composition (Figure S5F, Table S12) (n=6 males, mean age 30 ± 8.45; n=6 females, mean age 28 ± 11). These data were corrected for BMI due to significant differences observed in this dataset (Figure S5B-D). Next, we asked whether age-dependent decline in beta cell function (Figure 2F) also depended on sex. We divided our GSIS data into age groups similar to our single-cell RNAseq metadata analysis (Figures 1,2), namely adults (18–48 y.o.) and aged adults (≥ 50 y.o.). Strikingly, beta cells from adult men secreted almost two-times more insulin than women, but male beta cells suffered an age-dependent functional decline, whereas beta cell function in women was maintained throughout adulthood (Figure 5B). These analyses provide evidence that alpha and beta cell function, as well as their aging-related functional declines, are significantly different between men and women (Figure 5A-B).

**Figure 5.**
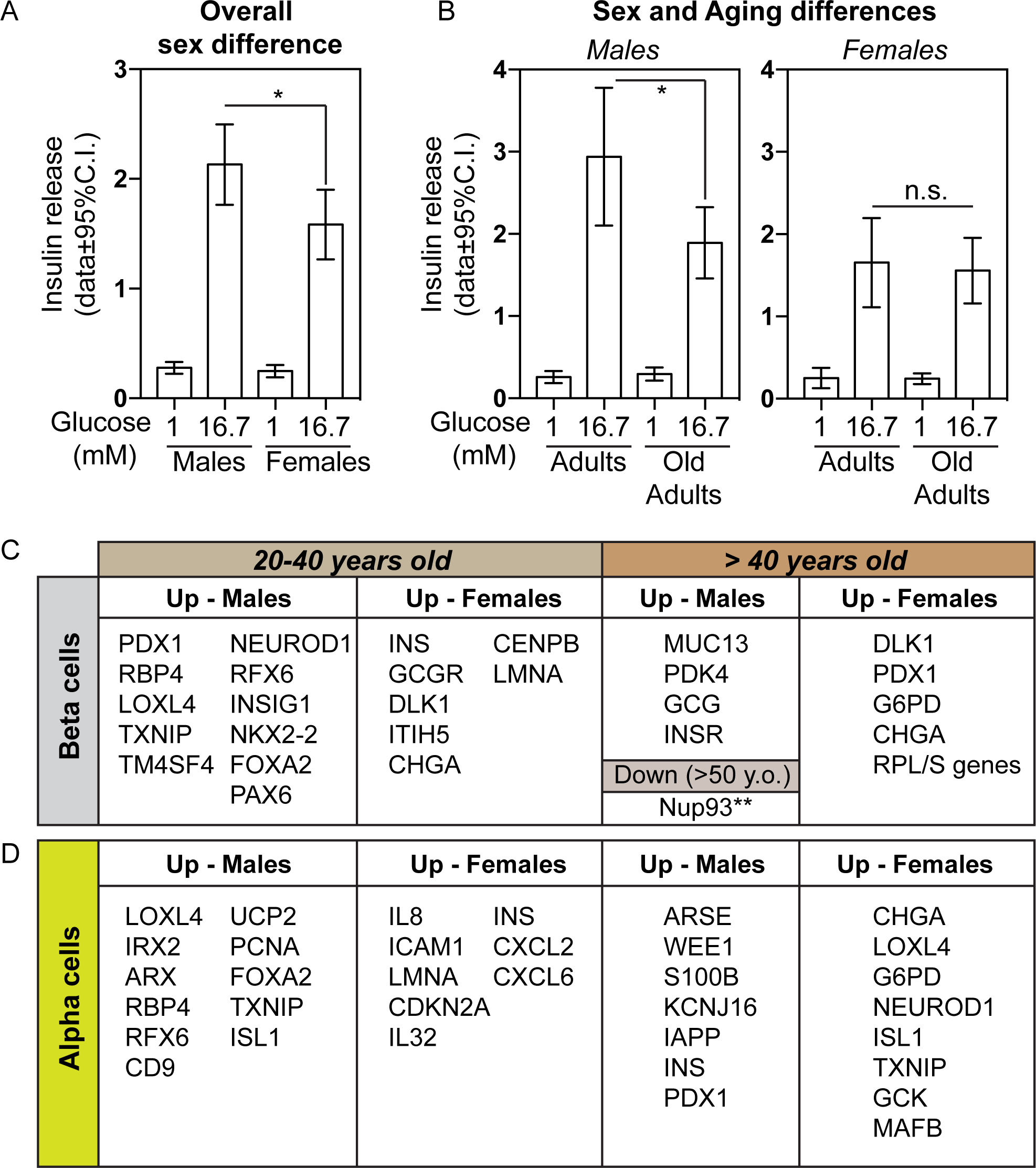
Effects of age and sex on islet cell function and transcriptional signature. Male beta cells secrete more insulin and suffer an age-related functional decline. **(A)** Glucose-stimulated insulin secretion (GSIS) assay performed on islets isolated from males and females (n=72 males, n=56 females). **(B)** GSIS assay data on islets isolated from males and females aged 18-to-48y.o. (n=30-37 males, n=15 females) and 50y.o. or older (n=34-38 males, n=37-41 females). Islets were exposed to 1mM and 16.7mM and insulin secretion is displayed as a relative % of the total islet insulin content. In (A), *p = 0.0164 and in (B), *p = 0.0136; n.s. = not significant. Error bars represent the 95% confidence interval of the data. In **(C)** and **(D)**, genes differentially expressed in beta and alpha cells, respectively, of men and women aged 20-to-40y.o. or 40.y.o. or older.

Next, we tested whether the observed age- and sex-dependent functional differences observed for both alpha and beta cells was associated with changes to their transcriptomes. We performed differential gene expression analysis of alpha and beta cells derived from N.D. men and women that were either 20–40 y.o. or ≥ 40 y.o. (Figure 5C-D). This age range was selected because our dataset only contained one female donor above 50 years of age. Nevertheless, young adult male beta cells expressed higher levels of genes associated with beta cell identity (e.g., PAX6, NKX2-2, PDX1), alpha cell identity (e.g., LOXL4, TM4SF4), and T2D (e.g., TXNIP) (Figure 5C). Similarly, male alpha cells expressed higher levels of genes involved in alpha cell identity (e.g., LOXL4, ARX), beta cell identity (FOXA2, CD9), and diabetes (e.g., TXNIP) (Figure 5D). Female beta cells, however, had higher levels of the beta cell marker genes, INS and DLK1, and the glucagon receptor (GCCR), among others (Figure 5C, Table S13). Aging affected male and female cells differently, with aged male alpha and beta cells exhibiting up-regulation of genes associated with other pancreatic cell types (e.g., ductal cells (MUC13)) (Figure 5C-D). Such age-related effects were not observed in female alpha or beta cells, but which instead had expressed high levels of genes involved in cell function, cell identity, and glucose metabolism (Figure 4C, Table S13). These results show that aging of alpha and beta cells is sex-specific and is associated with a loss of genes driving beta cell identity and function. This may explain the decline in beta cell function observed in males (Figure 5B), and why men suffer a more pronounced deterioration in glucose homeostasis during aging (Basu et al., 2006).

### Age-dependent breakdown of nuclear integrity in aged human beta cells

Beta cells are long-lived cells that become functional impaired with age (Figure 2F) (Arrojo e Drigo et al., 2019; Cnop et al., 2010; Perl et al., 2010; Westacott et al., 2017)). We previously established in rodents that long-lived cells contain long-lived proteins (Arrojo e Drigo et al., 2019; D’Angelo et al., 2009; Toyama et al., 2013). Most long-lived proteins localize to the nucleus, including a number of structural proteins, called nucleoporins, that are integral parts of the nuclear pore complex. In neurons, the lack of protein turnover in nuclear pore complexes leads to age-dependent loss of the nucleoporin, Nup93. This in turn leads to breakdown of the nuclear permeability barrier, and inappropriate exchange of material between the nucleus and cytoplasm (D’Angelo et al., 2009).

Our current analysis of human beta cells revealed that expression of the *NUP93* gene was significantly down-regulated with age in males (Figure 5C, Table S13, FDR<5%). To determine whether this phenotype extended to the protein level in humans, which have a much longer lifespan than rodents, we analyzed levels of pancreatic Nup93 protein levels in samples from young (< 30 y.o.) or aged (> 60 y.o.) N.D. donors from both sexes. First, we determined Nup93 and phenylalanine-glycine repeat (FG)-containing nucleoporins (FG-Nups) protein levels in snap-frozen sections. Pancreatic cells from a 74 y.o. donor had low levels of Nup93 (Nup93^Lo^, Figure 6A, S6A). Given the scarcity of tissue banks with a sizable archive of fresh-frozen pancreatic sections from N.D. young and aged humans, we next assessed levels of Nup93 in paraffin-embedded sections of isolated islets from young and aged donors (Table S14). We quantified the relative levels of Nup93 and FG-Nups in single human beta cells using confocal microscopy and an automated image processing protocol (Figure S6-7, methods). We observed an almost linear relationship between levels of Nup93 and FG-Nups in young human beta cells. In contrast, aged human beta cells had relatively less Nup93, resulting in a shift of the Nup93/FG-Nups correlation up-leftward towards the FG-Nups axis, resulting in loss of the linear correlation seen in young humans (r^2^ = 0.575 vs. r^2^ = 0.838, respectively, Figure 6B). Loss of Nup93 in aged rat neurons causes weakening of the nuclear permeability barrier and accumulation of cytoplasmic proteins in the nucleoplasm. For example, tubulin-beta-3 (Tubb3), which is normally cytoplasmic, forms intra-nuclear “rods” in aged neurons (D’Angelo et al., 2009). Accordingly, the frequency of Tubb3-positive intra-nuclear rods in beta cells from aged donors was 2-fold higher than in the younger cohort (Figure 6C). Together, these data provide the first evidence that aged human beta cells lose Nup93, and therefore exhibit impaired functionality of the nuclear permeability barrier.

**Figure 6.**
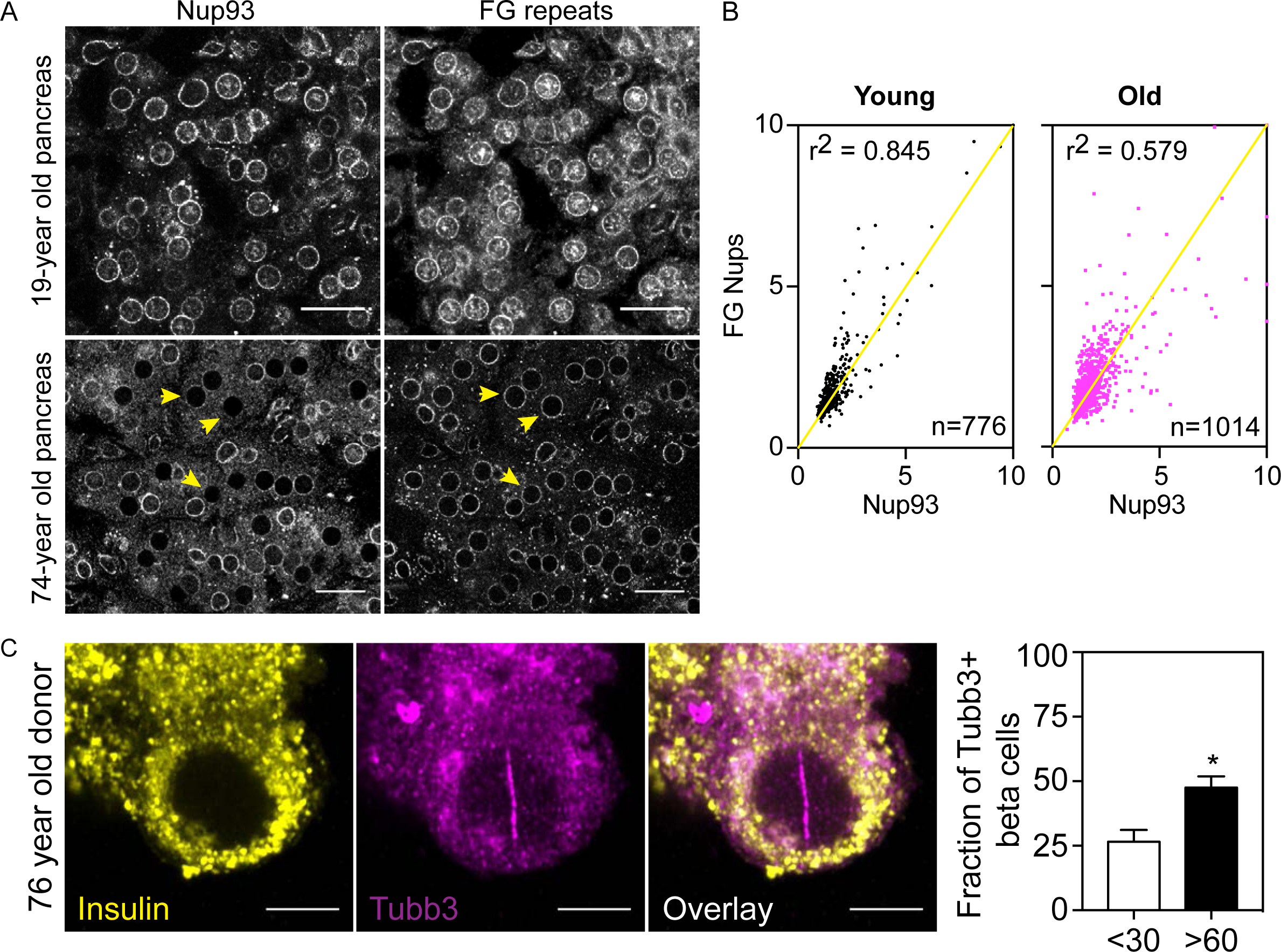
Age-dependent loss of LLP Nup93 in old human beta cells. **(A)** Confocal images of Nup93 and FG-Nups immunohistochemistry of frozen 19- or 74-year old pancreas. Yellow arrowheads show Nup93-low cells. **(B)** Relative levels of Nup93 and FG-Nups in beta cells in young and old humans. Yellow line represents a perfect linear correlation r^2^=1. Samples located above the yellow line are Nup93-low cells. **(C)** Confocal imaging of Tubb3 (magenta) in the beta cell (yellow) nucleus from a 76-year-old human. Right, percentage of Tubb3^+^ beta cells in young (<30 years old, n = 32 islets total, n = 5 donors) and old (>60 years old, n = 36 islets, n = 5 donors) subjects. **(D)** Illustration describing the age mosaicism at the NPC level due to heterogeneous turnover of LLPs in post-mitotic cells. Left, an NPC from a young cell where all Nups are of the same age and have an intact permeability barrier. Center, an NPC of an adult cell where few units of Nup93/133 (light gray) were replaced by new units (dark gray). Right, an NPC from an old post-mitotic cell and with a specific loss of Nup93 and a leaky permeability barrier. In (C), data shown as +/-standard error of the mean, *p < 0.01 by unpaired Student’s t-test. Scale bars, (A) 25 µm, (C) 5 µm.

## Discussion

Aging increases the risk of developing cardio-vascular and metabolic diseases, such as T2D. The islets of Langerhans contain mostly long-lived cells that persist throughout an organism’s lifetime, which in humans can be more than a century (Arrojo e Drigo et al., 2019; Perl et al., 2010). Beta cells derived from aged individuals or patients with T2D can display loss of cell identity and function (Enge et al., 2017; Guo et al., 2013; Talchai et al., 2012; Westacott et al., 2017). However, it has also been shown that aging correlates with changes in beta cell heterochromatin marks that reinforce the expression of beta cell identity genes and enhance insulin secretion (Avrahami et al., 2015), whereas another study found no age-related changes in beta cell function (Almaça et al., 2014). These contradictory results could be due to limitations in the age range and composition (percentage of males and females) of the islet donor cohort, or sample size (Almaça et al., 2014; Avrahami et al., 2015; Enge et al., 2017). These analyses are further complicated by the fact that beta cells are heterogeneous, with distinct transcriptional and functional profiles (Camunas-Soler et al., 2019; Dorrell et al., 2016; Johnston et al., 2016; van der Meulen et al., 2017; Van Schravendijk et al., 1992; Segerstolpe et al., 2016; Xin et al., 2018).

Here we describe the integration of human islet single-cell RNAseq datasets to determine the aging signature of major endocrine cell types in islets from mostly N.D. and T2D adult donors (Figure 1). Using this approach, we show that ∼75% of beta cells from aged N.D. donors had a transcriptional signature that was similar to T2D beta cells. This could explain declines in stimulated insulin release observed in islets from aged donors (Figure 2-3). Strikingly, a fraction of beta cells from both aged N.D. donors and age-matched T2D donors maintained a youthful transcriptional signature (16% and 5%, respectively) that was associated with high-functioning beta cells, including higher levels of expression of the incretin receptor, *GLP-1R* (Figure 2-3). In contrast, delta cells exhibited much smaller age-associated transcriptional shifts, and as a result we did not observe significant changes in delta cell sub-populations (Figures 2 and 4). However, in T2D islets, delta cells displayed a transcriptional signature associated with impairments in both exocytosis and glucose metabolism. This transcriptional shift was associated with an almost complete reconfiguration of delta cell sub-populations in T2D. Although the effects of T2D on human delta cells remain poorly characterized, these observations support previous observations that the structure-function of delta cells is altered in diabetic animal models (Abdel-Halim et al., 1993; Guardado Mendoza et al., 2015; Weir et al., 1981).

Men and women maintain glucose homeostasis in different ways and have different risk of developing T2D as they age (Basu et al., 2003, 2006; Logue et al., 2011). A recent study has described sex-specific differences in islet cells from N.D. individuals, including divergent patterns of DNA methylation, expression of islet genes, and insulin release (Hall et al., 2014). In addition, female islet cells released ∼20% more insulin than male islets, a difference that we did not detect (Figure 5). In fact, we found that islets from young and aged adult donor women secreted less insulin and glucagon than male islets, and were protected from age-related declines in beta cell function (Figure 5, S5). Furthermore, with the exception of elevated expression of the *RPS4X* gene in aged females (Table S13), we did not recapitulate the findings of Hall et al., 2014. This is likely due to differences in experimental setup (e.g., single-cell versus whole islet gene expression measurements).

Importantly, our analysis revealed sex-specific differences in the expression of cell identity genes and how they were impacted by aging. Genes associated with cell identity, islet cell function, and glucose metabolism were all down-regulated in cells from aged men, but not in aged women (Figure 5). These findings highlight the differential contribution of age and sex to the transcriptional and functional phenotypes of major cell types in the islets of Langerhans. In addition, our analyses provide detailed insights into the effects of aging on remarkably long-lived cell types, which are under lifelong pressure to sustain optimal levels of function to maintain glucose homeostasis for decades.

---

**Supplementary Figure 1 – Meta-analysis of single islet cell RNA seq datasets. (A)** and **(B)**, tSNE clustering of the single cell datasets analyzed in this study before and after batch correction with the Combat algorithm, respectively. **(B) (C)** Same as in (B), however the samples are labelled according to the health status of their respective donor, namely N.D.s (black dots) and T2D donors (blue dots). **(D)** to **(I)**, same as in (B-C), however cells are labelled by the normalized Z-score of pancreatic cell marker genes: *INS* for beta cells, *GCG* for alpha cells, *SST* for delta cells, *KRT19* for ductal cells, *PRSS1* for acinar cells and *PPY* for PP cells. **(J)** Relative contribution of the hormone genes *INS, GCG* and *SST* to the total transcriptomic load from single alpha, beta and delta cells. **(K)** Relative cell type composition of each donor dataset analyzed. **(L)** Venn diagrams representing the overlap in the number of genes between the three studies used in our meta-analysis approach. **(M)** Pie charts representing the fraction of genes that change with age in old alpha cells and beta cells. Genes differentially expressed that are associated with an alpha cell or beta cell signature. Light and dark green, alpha cell identity (alphaID) genes and in light and dark blue, beta cell identity genes (betaID) that are up or down-regulated in these cell types, respectively.

**Supplementary Figure 2 – Alpha, beta and delta cell sub-populations. (A)** Hierarchical clustering analysis with Pagoda2 of single beta cells highlighting genes that represent in a donor specific signature (21y.o. from (Enge et al., 2017)). Gene names are listed on top. **(B)** Panels of heatmaps showing the normalized expression of beta cell identity marker genes and beta cell heterogeneity markers described previously and in **(C)**, beta cell disallowed genes. Gene names are listed on top. **(D)** Same as (A), however for single alpha cells analyzed with Pagoda2. Heatmap highlights genes that represent alpha cell identity markers and donor specific signatures (21y.o. from (Enge et al., 2017)). Gene names are listed on top. **(E)** Identification of different types of delta cells in N.D. donors based on their transcriptional states by hierarchical clustering analysis using Pagoda2. Heatmaps show the normalized expression of cell identity marker genes (top panel, gene names listed on the right) and representative genes differentially expressed in two major delta cell state identified. Representative genes were chosen based on their Z-scores. **(F)** Relative distribution of delta cells into each state identified with Pagoda2 and as a function of age. Samples were divided into young (<6 y.o.), adults (20-to-40y.o.) and old adults (40-to-68y.o.).

**Supplementary Figure 3 – Pagoda2 and Monocle analysis of old and T2D beta cells.** Identification of different types of old and T2D beta cells based on their transcriptional states. In**(A)** Hierarchical clustering analysis of beta cells from old N.D. donors and T2D donors. Heatmaps show the normalized expression of representative genes differentially expressed in each beta cell state identified. Below each heatmap, pathway enrichment analysis (Metascape) for each cell state is shown. Representative genes were chosen based on their Z-scores. Black arrows indicate pathways that are positively regulated. For a full list of differentially expressed genes in each cell state, please refer to Table S10. **(B)** Relative distribution of beta cells from N.D. and T2D donors in both states shown in (A). **(C)** Relative donor composition of each beta cell state shown in (A). **(D)** Pseudotime ordering of single old adult and T2D beta cells using Monocle (v2.4). Each beta cell is shown as a single dot and beta cells states identified with Monocle are shown in different colors. **(E)** Relative distribution of beta cells from N.D. and T2D donors in the Monocle beta cell states shown in (D). **(F)** Same as (D), however each single beta cell is colored by the age of their respective islet donor. **(G-K)** 2-D pseudotime jitter plots displaying the expression pattern of representative genes differentially expressed in beta cell states identified with Monocle. The black line in bold indicates the mean expression pattern across the pseudotime tree. Each cell is represented by a dot and each dot is colored by their respective cell state.

**Supplementary Figure 4 – Analysis of delta cell single cell data with Monocle v2.4. (A)** Pseudotime ordering of single old adult and T2D delta cells using Monocle (v2.4). Each delta cell is shown as a single dot. Here, each single delta cell is colored by the age of their respective islet donor. **(B)** Relative distribution of delta cells from N.D. and T2D donors in each cell states shown in (A).

**Supplementary Figure 5 – Structure-function data of islets from men and women. (A)** Insulin content of male and female islets, assayed at the end of each GSIS experiment. GSIS data is shown in Figure 4A. In **(B)** and **(C)**, data from islet perifusion experiments available from the HPAP consortium database. Curves from males and females are shown in black and red lines, respectively. **(D)** Body mass index (BMI) for samples shown in (B-D). **(E)** Oxygen consumption rate (OCR) of isolated islets from males and females (represented by black and red lines, respectively). Data available from the HPAP consortium.

**Supplementary Figure 6 -** Immunohistochemistry of human pancreas. (A) Immunostaining of frozen human pancreas from a 74-year old cadaveric donor with anti-Mab414 (white) and - Nup93 (magenta) antibodies. Inset on the right shows a close up on two cells indicated by the yellow box. (B) Immunohistochemistry of paraffin-embedded isolated human islets from an 18-year old and a 60-year old donor with anti-Mab414, -Nup93 and -insulin antibodies. Insets show a close up of cells inside the dashed yellow box. Arrowheads indicate different beta cells with varying levels of Nup93. Scale bars 25um, in insets 5um.

**Supplementary Figure 7 -** Work-flow of the image processing and total intensity calculation for Nup93 and Mab414 (FG-Nups) at NPCs. The DAPI channel image was used for image segmentation. Next, a binary mask was created from the segmented image which was later used to find the rim around nucleus and calculate the relative antibody signal at the NPC region and the background signal in both Nup93 and Mab414 channel images.

**Supplementary Table 1 – Donor metadata of single cell RNAseq datasets**

**Supplementary Table 2 – Islet cell marker gene expression**

**Supplementary Table 3 – Genes differentially expressed in adult and old beta cells**

**Supplementary Table 4 – Genes differentially expressed in adult and old alpha cells**

**Supplementary Table 5 – Genes differentially expressed in the beta cell states identified with Pagoda2**

**Supplementary Table 6 – Genes differentially expressed in the alpha cell states identified with Pagoda2**

**Supplementary Table 7 – Relative composition of alpha, beta and delta cell states in young, adults and old adults.**

**Supplementary Table 8 – Genes differentially expressed in the delta cell states identified with Pagoda2**

**Supplementary Table 9 – GSIS data from isolated human islets and donor profile**

**Supplementary Table 10 - Genes differentially expressed in beta cells in old N.D. and T2D donors identified with Pagoda2**

**Supplementary Table 11 - Genes differentially expressed in beta cell states identified with Monocle v2.4**

**Supplementary Table 12 - Genes differentially expressed in delta cell states identified with Monocle v2.4**

**Supplementary Table 13 - Genes differentially expressed in beta cells from males and females**

**Supplementary Table 14 - Genes differentially expressed in alpha cells from males and females**

**Supplementary Table 15 – Alpha and beta cell composition from the HPAP consortia dataset**

## Acknowledgements

This work was supported by grants to M.H from the National Institutes of Health Transformative Research Award grant R01 NS096786, the Keck Foundation and the NOMIS Foundation and by a grant from the National Institutes of Health and Human Islet Research Network (1U01DK120447-01) to R.AeD, P.E.M, and M.H. We thank the organ procurement organizations across Canada, particularly the Human Organ Procurement and Exchange (HOPE) program in Edmonton and the Trillium Gift of Life Network (TGLN) in Ontario, for their work in obtaining human pancreas for research. Conventional light microcopy work utilized the Waitt Advanced Biophotonics Core Facility of the Salk Institute with funding from NIH-NCI CCSG: P30 014195, NINDS Neuroscience Core Grant: NS072031 and the Waitt Foundation. Bioinformatics leveraged the Next Generation Sequencing Core Facility of the Salk Institute with funding from NIH-NCI CCSG: P30 014195, the Chapman Foundation and the Helmsley Charitable Trust, and Max Shokhirev and Max Chang of The Razavi Newman Integrative Genomics and Bioinformatics Core Facility of the Salk Institute with funding from NIH-NCI CCSG: P30 014195, and the Helmsley Trust. R.AeD is supported by an American Diabetes Association postdoctoral fellowship (#1-18-PMF-007). The authors are also thankful to David O’Keefe (Salk Institute) for critical inputs on manuscript editing and revision. Finally, we are indebted to organ donors and their families for their generous support of scientific research.

## References

Abdel-Halim, S.M., Guenifi, A., Efendic, S., and Ostenson, C.-G (1993). Both somatostatin and insulin responses to glucose are impaired in the perfused pancreas of the spontaneously noninsulin-dependent diabetic GK (Goto-Kakizaki) rats. Acta Physiol. Scand. 148, 219–226.

Aguayo-Mazzucato, C., van Haaren, M., Mruk, M., Lee, T.B., Crawford, C., Hollister-Lock, J., Sullivan, B.A., Johnson, J.W., Ebrahimi, A., Dreyfuss, J.M., et al. (2017). Beta Cell Aging Markers Have Heterogeneous Distribution and Are Induced by Insulin Resistance. Cell Metab. 25.

Almaça, J., Molina, J., Arrojoe Drigo, R., Abdulreda, M.H., Jeon, W.B., Berggren, P.-O., Caicedo, A., and Nam, H.G. (2014). Young capillary vessels rejuvenate aged pancreatic islets. Proc. Natl. Acad. Sci. 111, 17612–17617.

Anderwald, C., Gastaldelli, A., Tura, A., Krebs, M., Promintzer-Schifferl, M., Kautzky-Willer, A., Stadler, M., DeFronzo, R.A., Pacini, G., and Bischof, M.G. (2011). Mechanism and effects of glucose absorption during an oral glucose tolerance test among females and males. J. Clin. Endocrinol. Metab.

Arda, H.E., Li, L., Tsai, J., Torre, E.A., Rosli, Y., Peiris, H., Spitale, R.C., Dai, C., Gu, X., Qu, K., et al. (2016). Age-dependent pancreatic gene regulation reveals mechanisms governing human β cell function. Cell Metab. 23.

Arrojoe Drigo, R., Ali, Y., Diez, J., Srinivasan, D.K., Berggren, P.O., and Boehm, B.O. (2015). New insights into the architecture of the islet of Langerhans: a focused cross-species assessment. Diabetologia 58, 2218–2228.

Arrojoe Drigo, R., Lev-Ram, V., Tyagi, S., Ramachandra, R., Deerinck, T., Bushong, E., Phan, S., Orphan, V., Lechene, C., Ellisman, M.H., et al. (2019). Age mosaicism across multiple scales in adult tissues. Cell Metab.

Avrahami, D., Li, C., Zhang, J., Schug, J., Avrahami, R., Rao, S., Stadler, M.B., Burger, L., Schübeler, D., Glaser, B., et al. (2015). Aging-dependent demethylation of regulatory elements correlates with chromatin state and improved β cell function. Cell Metab.

Basu, R., Breda, E., Oberg, A., Powell, C., Dalla Man, C., Basu, A., Vittone, J., Klee, G., Arora, P., Jensen, M., et al. (2003). Mechanisms of the Age-Associated Deterioration in Glucose Tolerance. Diabetes 52, 1738–1748.

Basu, R., Man, C.D., Campioni, M., Basu, A., Klee, G., Toffolo, G., Cobelli, C., and Rizza, R.A. (2006). Effects of age and sex on postprandial glucose metabolism differences in glucose turnover, insulin secretion, insulin action, and hepatic insulin extraction. Diabetes 55, 2001–2014.

Buchwalter, A., and Hetzer, M.W. (2017). Nucleolar expansion and elevated protein translation in premature aging. Nat. Commun.

Cabrera, O., Berman, D.M., Kenyon, N.S., Ricordi, C., Berggren, P.-O., and Caicedo, A. (2006). The unique cytoarchitecture of human pancreatic islets has implications for islet cell function. Proc. Natl. Acad. Sci. U. S. A. 103, 2334–2339.

Camunas-Soler, J., Dai, X., Hang, Y., Bautista, A., Lyon, J., Suzuki, K., Kim, S.K., Quake, S.R., and MacDonald, P.E. (2019). Pancreas patch-seq links physiologic dysfunction in diabetes to single-cell transcriptomic phenotypes. BioRxiv.

Cnop, M., Hughes, S.J., Igoillo-Esteve, M., Hoppa, M.B., Sayyed, F., Van De Laar, L., Gunter, J.H., De Koning, E.J.P., Walls, G. V., Gray, D.W.G., et al. (2010). The long lifespan and low turnover of human islet beta cells estimated by mathematical modeling of lipofuscin accumulation. Diabetologia 53, 321–330.

D’Angelo, M.A., Raices, M., Panowski, S.H., and Hetzer, M.W. (2009). Age-Dependent Deterioration of Nuclear Pore Complexes Causes a Loss of Nuclear Integrity in Postmitotic Cells. Cell 136, 284–295.

Doria, A., Patti, M.-E., and Kahn, C.R. (2008). The emerging genetic architecture of type 2 diabetes. Cell Metab. 8, 186–200.

Dorrell, C., Schug, J., Canaday, P.S., Russ, H.A., Tarlow, B.D., Grompe, M.T., Horton, T., Hebrok, M., Streeter, P.R., Kaestner, K.H., et al. (2016). Human islets contain four distinct subtypes of β cells. Nat. Commun. 7, 11756.

Enge, M., Arda, H.E., Mignardi, M., Beausang, J., Bottino, R., Kim, S.K., and Quake, S.R. (2017). Single-Cell Analysis of Human Pancreas Reveals Transcriptional Signatures of Aging and Somatic Mutation Patterns. Cell.

Færch, K., Borch-Johnsen, K., Vaag, A., Jørgensen, T., and Witte, D.R. (2010). Sex differences in glucose levels: A consequence of physiology or methodological convenience? the Inter99 study. Diabetologia.

Fan, J., Salathia, N., Liu, R., Kaeser, G.E., Yung, Y.C., Herman, J.L., Kaper, F., Fan, J.B., Zhang, K., Chun, J., et al. (2016). Characterizing transcriptional heterogeneity through pathway and gene set overdispersion analysis. Nat. Methods.

Gannon, M., Kulkarni, R.N., Tse, H.M., and Mauvais-Jarvis, F. (2018). Sex differences underlying pancreatic islet biology and its dysfunction. Mol. Metab.

Guardado Mendoza, R., Perego, C., Finzi, G., La Rosa, S., Capella, C., Jimenez-Ceja, L.M., Velloso, L.A., Saad, M.J.A., Sessa, F., Bertuzzi, F., et al. (2015). Delta cell death in the islet of Langerhans and the progression from normal glucose tolerance to type 2 diabetes in nonhuman primates (baboon, Papio hamadryas). Diabetologia 58, 1814–1826.

Guo, S., Dai, C., Guo, M., Taylor, B., Harmon, J.S., Sander, M., Robertson, R.P., Powers, A.C., and Stein, R. (2013). Inactivation of specific β cell transcription factors in type 2 diabetes. J. Clin. Invest.

Gutierrez, G.D., Xin, Y., Okamoto, H., Kim, J., Lee, A.H., Ni, M., Adler, C., Yancopoulos, G.D., Murphy, A.J., and Gromada, J. (2018). Gene signature of proliferating human pancreatic a cells. Endocrinology.

Gutiérrez, G.D., Bender, A.S., Cirulli, V., Mastracci, T.L., Kelly, S.M., Tsirigos, A., Kaestner, K.H., and Sussel, L. (2017). Pancreatic β cell identity requires continual repression of non-β cell programs. J. Clin. Invest. 127.

Hall, E., Volkov, P., Dayeh, T., Esguerra, J.L. o. S., Salö, S., Eliasson, L., Rönn, T., Bacos, K., and Ling, C. (2014). Sex differences in the genome-wide DNA methylation pattern and impact on gene expression, microRNA levels and insulin secretion in human pancreatic islets. Genome Biol.

Hauge-Evans, A.C., King, A.J., Carmignac, D., Richardson, C.C., Robinson, I.C.A.F., Low, M.J., Christie, M.R., Persaud, S.J., and Jones, P.M. (2009). Somatostatin secreted by islet delta-cells fulfills multiple roles as a paracrine regulator of islet function. Diabetes 58, 403–411.

Helman, A., Klochendler, A., Azazmeh, N., Gabai, Y., Horwitz, E., Anzi, S., Swisa, A., Condiotti, R., Granit, R.Z., Nevo, Y., et al. (2016). p16(Ink4a)-induced senescence of pancreatic beta cells enhances insulin secretion. Nat. Med. 22, 412–420.

https://hpap.pmacs.upenn.edu/ Human Pancreas Analysis Program (HPAP) Database, a consortium under the Human Islet Research Network (RRID:SCR_014393).

Johnston, N.R., Mitchell, R.K., Haythorne, E., Pessoa, M.P., Semplici, F., Ferrer, J., Piemonti, L., Marchetti, P., Bugliani, M., Bosco, D., et al. (2016). Beta Cell Hubs Dictate Pancreatic Islet Responses to Glucose. Cell Metab. 24, 389–401.

Kalyani, R.R., Golden, S.H., and Cefalu, W.T. (2017). Diabetes and Aging: Unique Considerations and Goals of Care. Diabetes Care.

Logue, J., Walker, J.J., Colhoun, H.M., Leese, G.P., Lindsay, R.S., McKnight, J.A., Morris, A.D., Pearson, D.W., Petrie, J.R., Philip, S., et al. (2011). Do men develop type 2 diabetes at lower body mass indices than women? Diabetologia.

Mauvais-Jarvis, F., Manson, J.A.E., Stevenson, J.C., and Fonseca, V.A. (2017). Menopausal hormone therapy and type 2 diabetes prevention: Evidence, mechanisms, and clinical implications. Endocr. Rev.

van der Meulen, T., Mawla, A.M., DiGruccio, M.R., Adams, M.W., Nies, V., Dólleman, S., Liu, S., Ackermann, A.M., Cáceres, E., Hunter, A.E., et al. (2017). Virgin Beta Cells Persist throughout Life at a Neogenic Niche within Pancreatic Islets. Cell Metab. 25.

Ori, A., Toyama, B.H., Harris, M.S., Bock, T., Iskar, M., Bork, P., Ingolia, N.T., Hetzer, M.W., and Beck, M. (2015). Integrated Transcriptome and Proteome Analyses Reveal Organ-Specific Proteome Deterioration in Old Rats. Cell Syst. 1, 224–237.

Perl, S.Y., Kushner, J.A., Buchholz, B.A., Meeker, A.K., Stein, G.M., Hsieh, M., Kirby, M., Pechhold, S., Liu, E.H., Harlan, D.M., et al. (2010). Significant human β-cell turnover is limited to the first three decades of life as determined by in vivo thymidine analog incorporation and radiocarbon dating. J. Clin. Endocrinol. Metab.

Petersen, K.F., Befroy, D., Dufour, S., Dziura, J., Ariyan, C., Rothman, D.L., DiPietro, L., Cline, G.W., and Shulman, G.I. (2003). Mitochondrial dysfunction in the elderly: possible role in insulin resistance. Science.

Pullen, T.J., Khan, A.M., Barton, G., Butcher, S.A., Sun, G., and Rutter, G.A. (2010). Identification of genes selectively disallowed in the pancreatic islet. Islets.

Qiu, X., Mao, Q., Tang, Y., Wang, L., Chawla, R., Pliner, H.A., and Trapnell, C. (2017). Reversed graph embedding resolves complex single-cell trajectories. Nat. Methods.

Van Schravendijk, C.F., Kiekens, R., and Pipeleers, D.G. (1992). Pancreatic beta cell heterogeneity in glucose-induced insulin secretion. J. Biol. Chem. 267, 21344–21348.

Segerstolpe, A., Palasantza, A., Eliasson, P., Andersson, E.-M., Andreasson, A.-C., Sun, X., Picelli, S., Sabirsh, A., Clausen, M., Bjursell, M.K., et al. (2016). Single-Cell Transcriptome Profiling of Human Pancreatic Islets in Health and Type 2 Diabetes. Cell Metab. 24, 593–607.

Sicree, R.A., Zimmet, P.Z., Dunstan, D.W., Cameron, A.J., Welborn, T.A., and Shaw, J.E. (2008). Differences in height explain gender differences in the response to the oral glucose tolerance test - The AusDiab study. Diabet. Med.

Talchai, C., Xuan, S., Lin, H. V., Sussel, L., and Accili, D. (2012). Pancreatic β cell dedifferentiation as a mechanism of diabetic β cell failure. Cell 150, 1223–1234.

Taylor, R.C., and Dillin, A. (2011). Aging as an event of proteostasis collapse. Cold Spring Harb. Perspect. Biol. 3, 1–17.

Toyama, B.H., Savas, J.N., Park, S.K., Harris, M.S., Ingolia, N.T., Yates, J.R., and Hetzer, M.W. (2013). Identification of long-lived proteins reveals exceptional stability of essential cellular structures. Cell 154, 971–982.

Unger, R.H., and Orci, L. (2010). Paracrinology of islets and the paracrinopathy of diabetes. Proc. Natl. Acad. Sci. U. S. A. 107, 16009–16012.

Wang, Y.J., Golson, M.L., Schug, J., Traum, D., Liu, C., Vivek, K., Dorrell, C., Naji, A., Powers, A.C., Chang, K.M., et al. (2016). Single-Cell Mass Cytometry Analysis of the Human Endocrine Pancreas. Cell Metab.

Weir, G.C., Clore, E.T., Zmachinski, C.J., and Bonner-Weir, S. (1981). Islet secretion in a new experimental model for non-insulin-dependent diabetes. Diabetes 30, 590–595.

Westacott, M.J., Farnsworth, N.L., St Clair, J.R., Poffenberger, G., Heintz, A., Ludin, N.W., Hart, N.J., Powers, A.C., and Benninger, R.K.P. (2017). Age-dependent decline in the coordinated [Ca2+] and insulin secretory dynamics in human pancreatic islets. Diabetes 66, 2436–2445.

Xin, Y., Kim, J., Okamoto, H., Ni, M., Wei, Y., Adler, C., Murphy, A.J., Yancopoulos, G.D., Lin, C., and Gromada, J. (2016). RNA Sequencing of Single Human Islet Cells Reveals Type 2 Diabetes Genes. Cell Metab.

Xin, Y., Gutierrez, G.D., Okamoto, H., Kim, J., Lee, A.H., Adler, C., Ni, M., Yancopoulos, G.D., Murphy, A.J., and Gromada, J. (2018). Pseudotime ordering of single human B-cells reveals states of insulin production and unfolded protein response. Diabetes.

