## Supplementary figures and images for "Aging of human endocrine pancreatic cell types is heterogeneous and sex-specific"

### Supplementary Figures

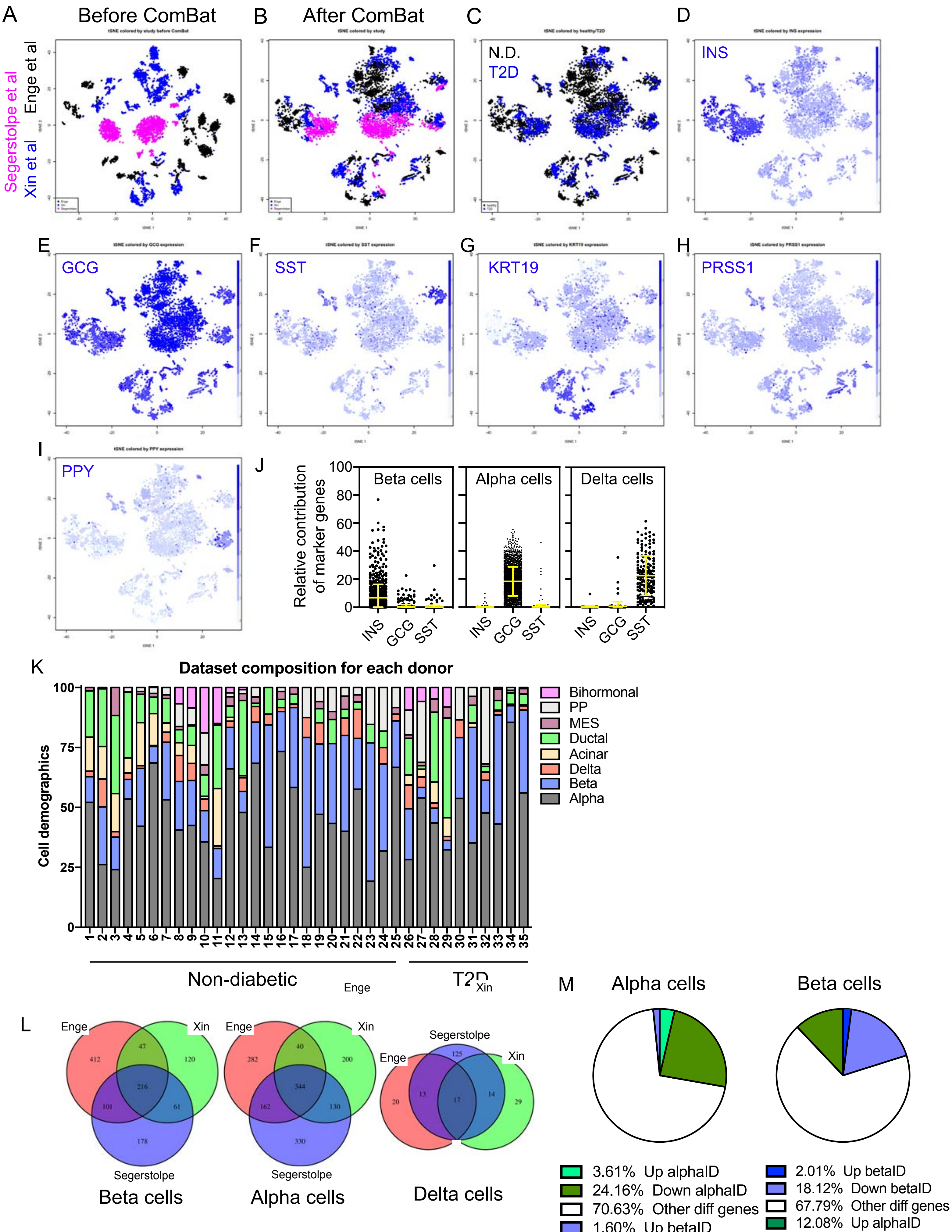

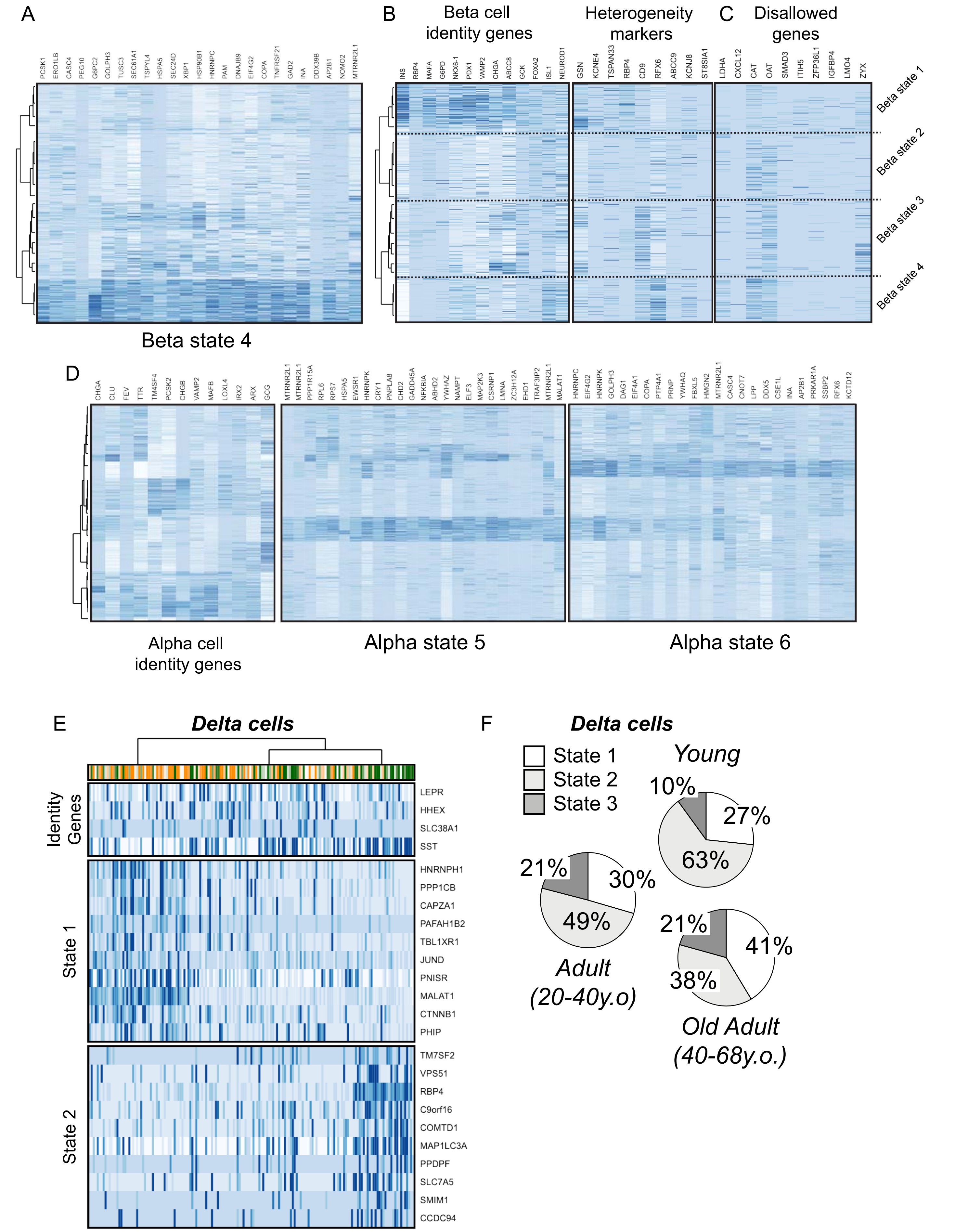

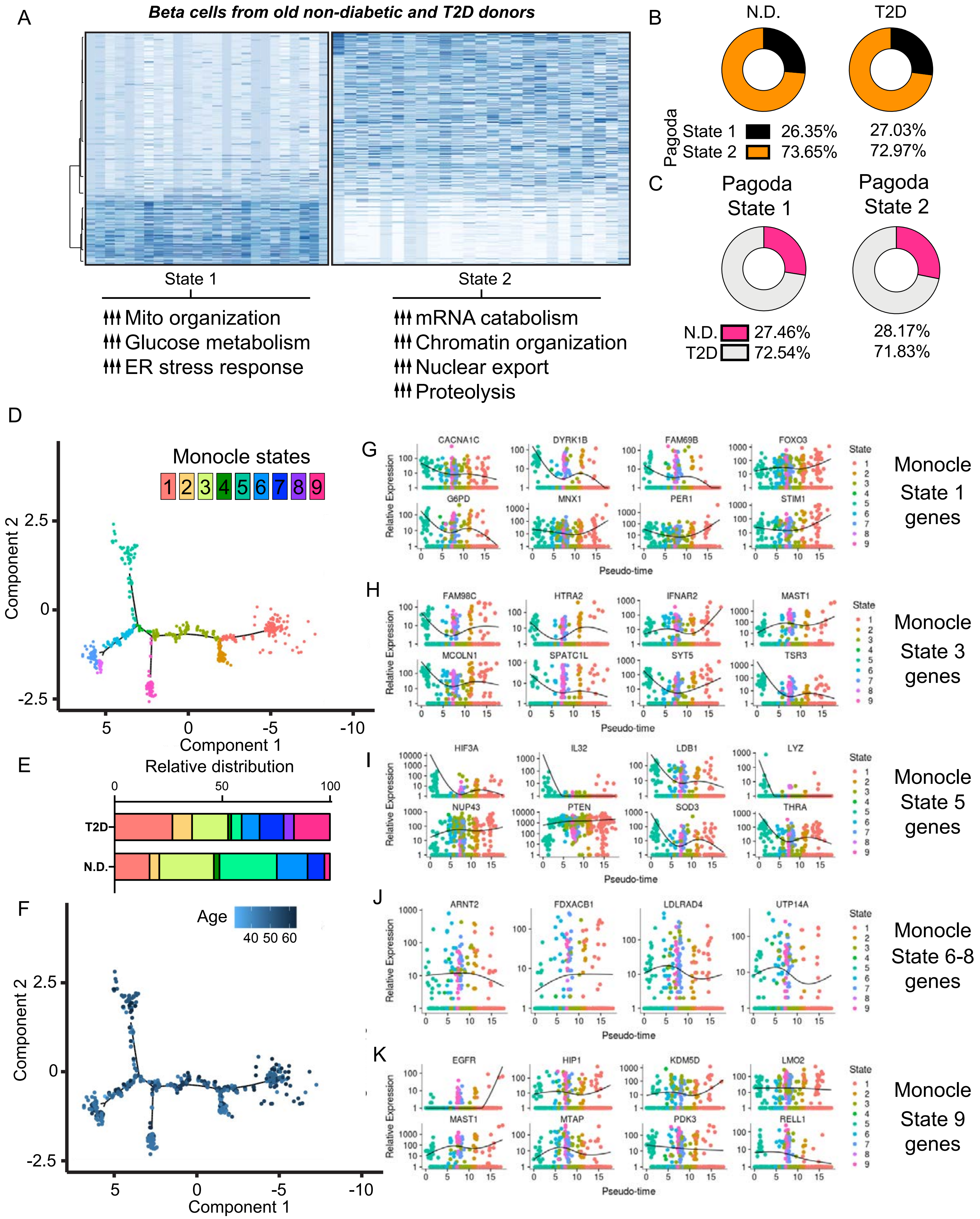

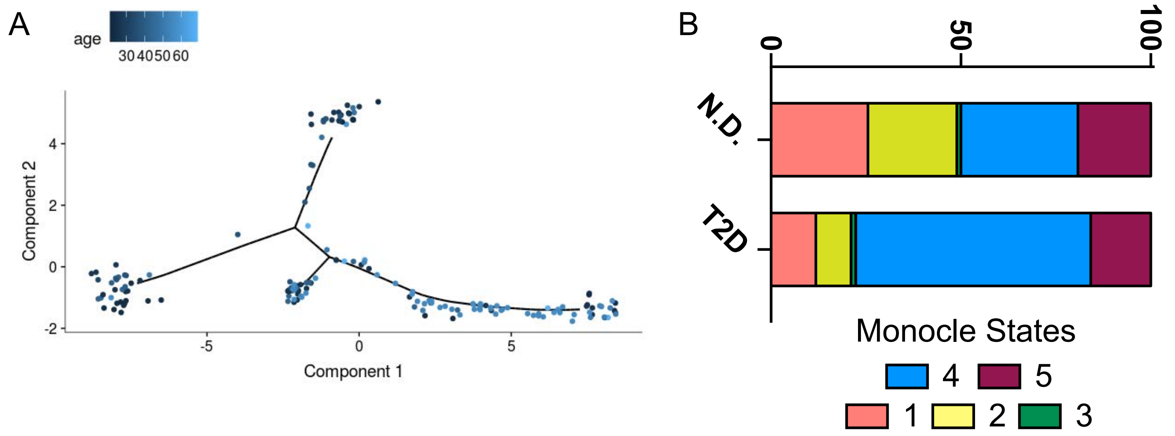

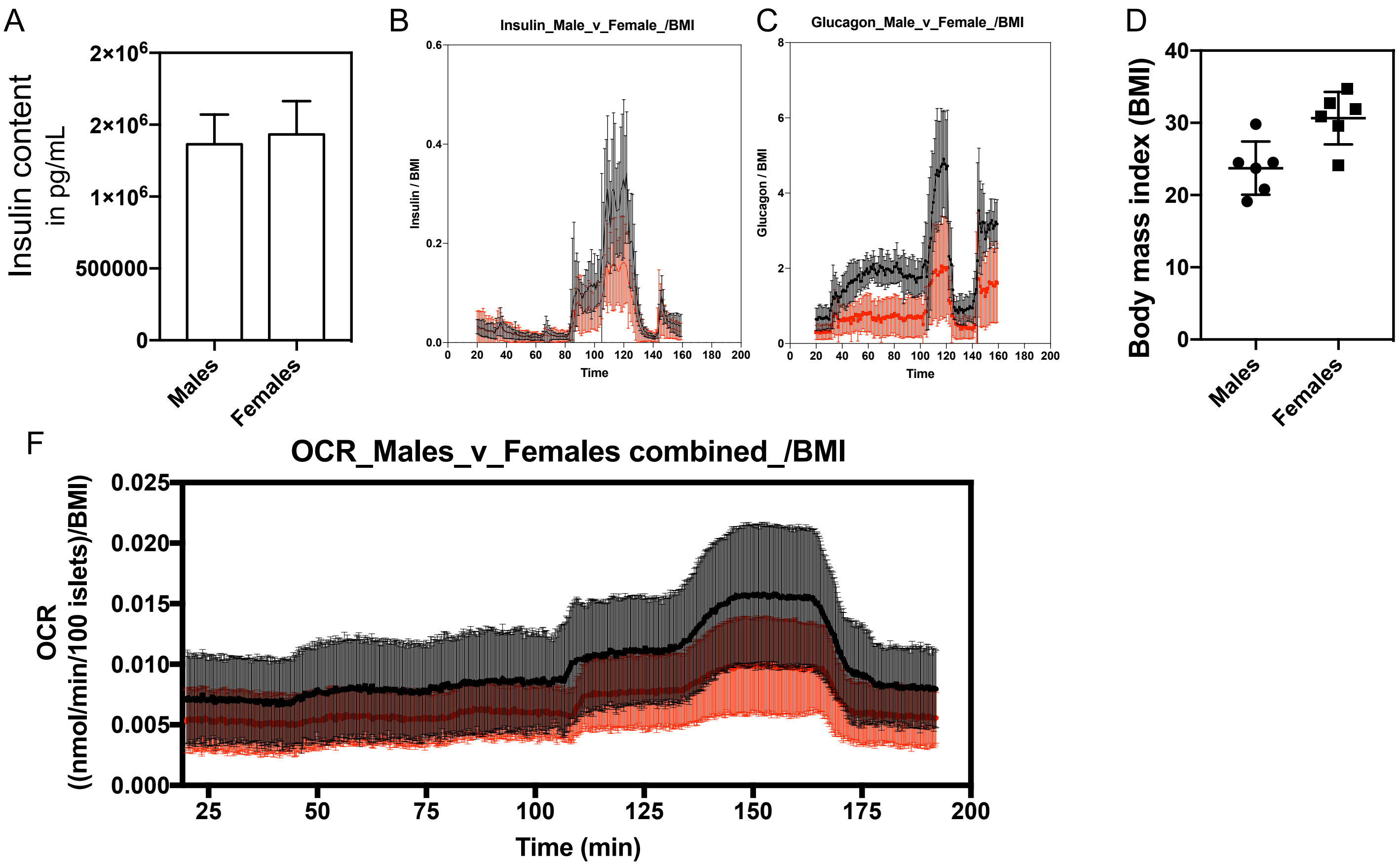

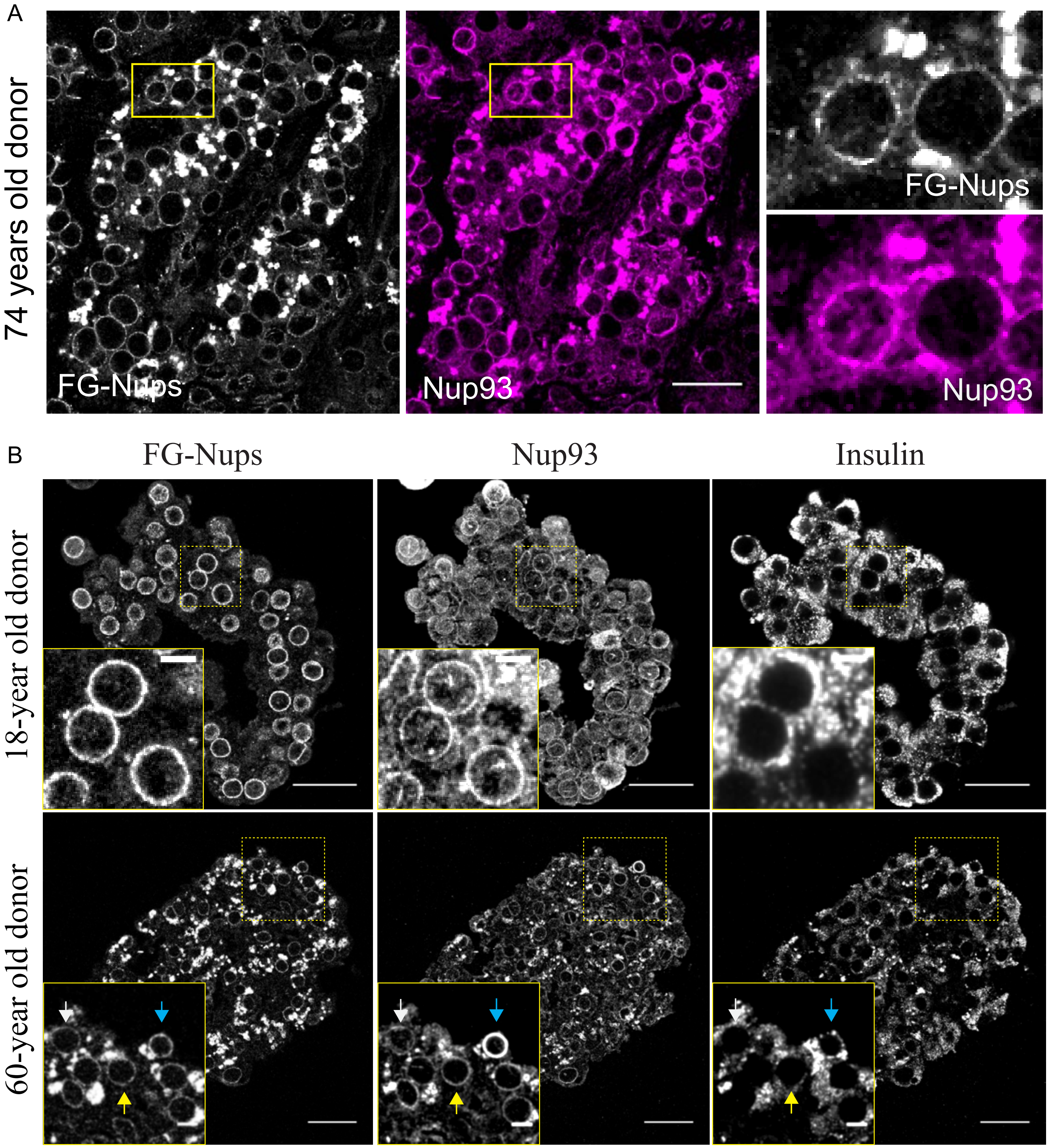

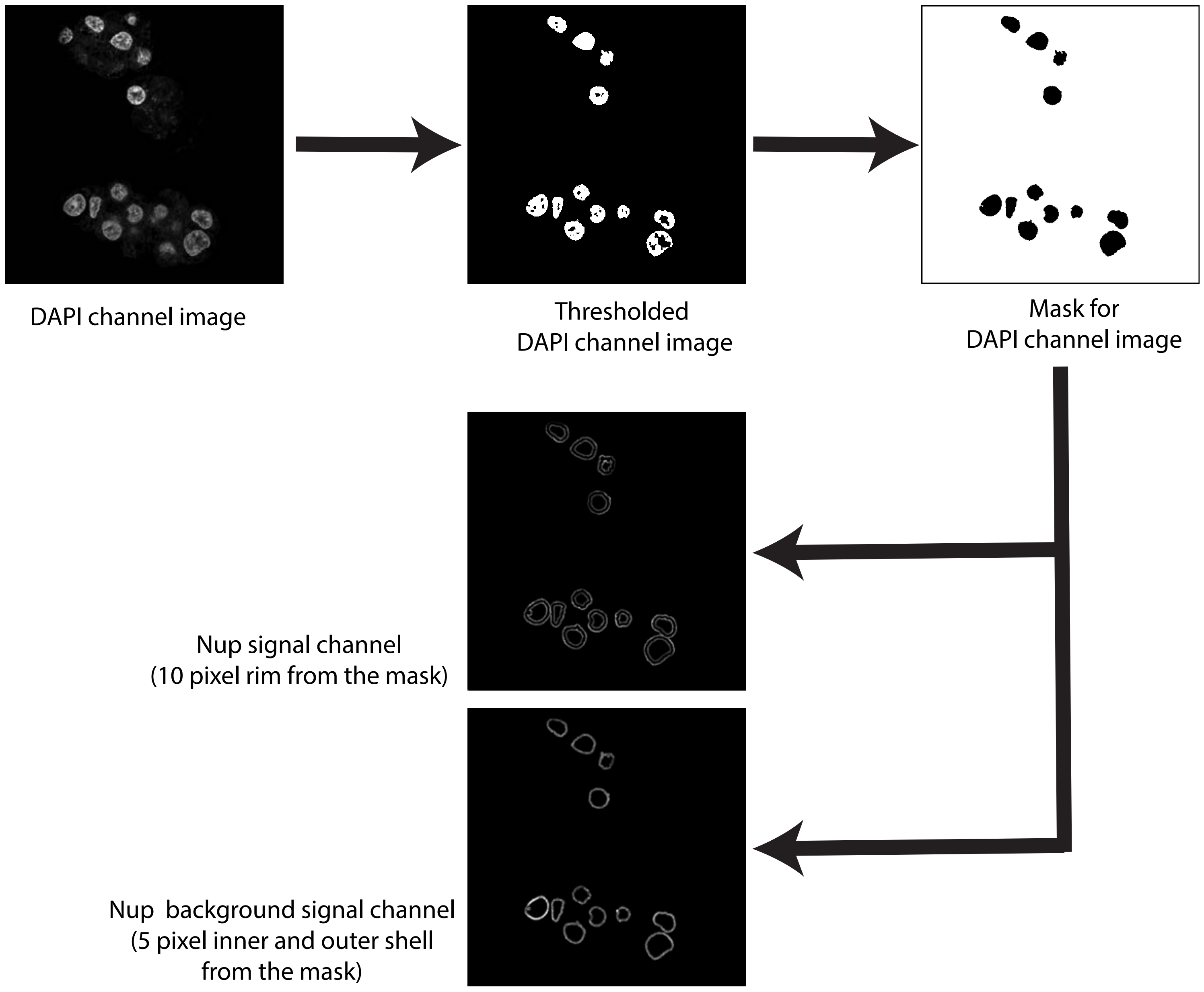
